# Sequence-based models predict TF binding with negligible contributions from motif cooperativity

**DOI:** 10.64898/2026.11.16.766980

**Authors:** Vladimir Mindel, Naama Barkai

## Abstract

How transcription factors (TFs) selectively recognize their genomic targets among numerous sites containing similar DNA motifs remains an open question. Cooperative binding at neighboring motif sites is widely considered a key mechanism of TF specificity, yet its genome-wide contribution remains unclear. Machine learning models can now predict TF binding directly from DNA sequence, providing opportunities to investigate its underlying determinants. Here, we leveraged BPNet models trained on ∼2,000 ENCODE TF binding profiles to systematically assess motif cooperativity. Although detected at individual loci, cooperativity made a negligible contribution to genome-wide binding predictions. Nevertheless, the models captured differences in binding preferences between TFs with similar motif preferences, even within the same cellular environment. These findings highlight the need to critically assess prevailing assumptions and explore alternative mechanisms underlying selective TF binding.

---

A fundamental question in gene regulation is how transcription factors (TFs) select their genomic binding sites. DNA-binding domains (DBDs) recognize short DNA sequence motifs, yet TFs typically occupy only a fraction of the many genomic sites containing these motifs^1,2^ (Fig. 1A-B). Moreover, TFs with similar or even identical motif preferences can occupy distinct sets of genomic sites. While DBD–motif recognition is relatively well understood, what determines the selective occupancy of individual motif sites remains poorly explained. Prevailing models attribute specificity primarily to chromatin accessibility and cooperative interactions with other TF^1,3-6^ (Fig. 1C). Yet, the capacity of these models to explain TF binding preferences has not been systematically assessed at the genomic scale.

**Figure 1.**
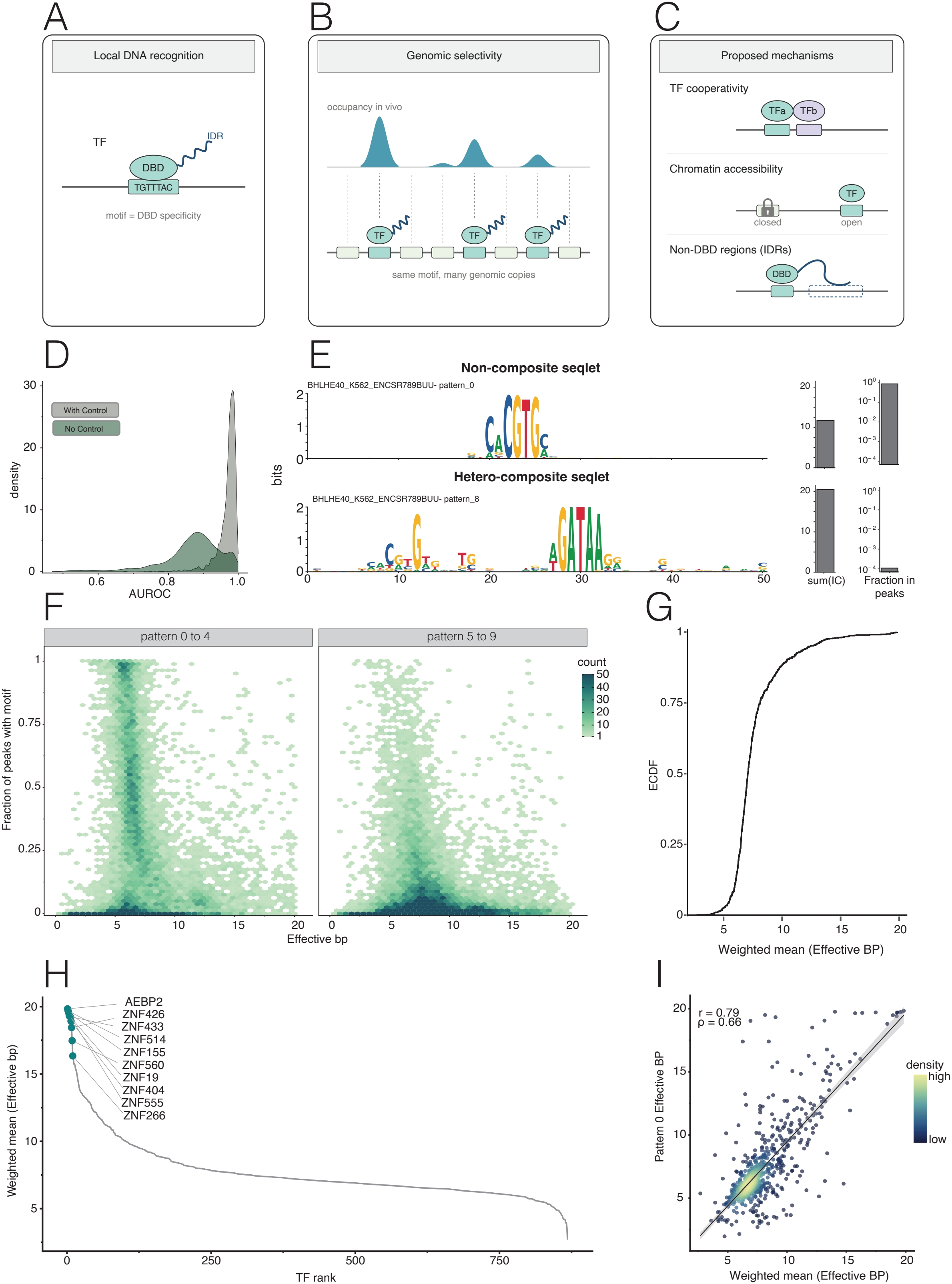
Composite motifs and high-information-content sequences are rare among seqlets identified as contributors to model-predicted TF binding. **(A–C)** Schematic overview of the problem of transcription factor (TF) binding specificity. (A-B) The DNA-binding domain (DBD) of a TF recognizes a specific DNA motif, while other protein regions, many of which are intrinsically disordered regions (IDRs), contribute to the selection of motif occurrences in genomes. (C) Prevailing models of TF binding specificity, including cooperative interactions between TFs and differential chromatin accessibility. **(D)** Distribution of BPNet model prediction performance (AUROC) across TFs, comparing predictions obtained with and without the free-DNA control tracks as an input. **(E)** Examples of sequence patterns (seqlets) identified by the BPNet interpretability framework for BHLHE40 TF in K562 cells. A non-composite seqlet (pattern 0, top) and a hetero-composite seqlet (pattern 8, bottom) are shown as contribution weight matrices (CWMs), together with their total information content (IC) and fraction of peaks containing the pattern. **(F)** Relationship between the effective sequence length (effective bp) of individual seqlets and the fraction of binding peaks containing a motif, shown separately for patterns 0–4 (left) and patterns 5–9 (right). Point density represents the number of observations. **(G)** Cumulative distribution of the weighted mean effective sequence length across peaks in all BPNet TF models. **(H)** Weighted mean effective sequence length across TF models, ranked from highest to lowest. The top-ranked TFs are indicated. **(I)** Relationship between the effective sequence length of the dominant pattern (pattern 0) and the weighted mean effective sequence length across all patterns. Each point represents a TF model. Pearson correlation (r) and Spearman correlation (ρ) are indicated.

Recent advances in applying machine learning to functional genomic data offer a powerful approach to investigating TF binding specificity. Deep learning models such as BPNet^7^ and AlphaGenome^8^ can predict TF binding profiles from genomic DNA sequences. BPNet was recently applied across more than 2,000 ENCODE TF ChIP-seq experiments (ENCODE Project Consortium^9^), demonstrating high accuracy in predicting binding profiles of hundreds of TFs (Fig. 1D, Fig.S1). The models therefore capture sequence information relevant to genomic specificity, providing an opportunity to systematically evaluate the contributions of proposed specificity mechanisms.

We leveraged BPNet models trained on ENCODE data to examine the sequence determinants underlying their binding prediction^7,10,11^. The models’ interpretation framework identifies sequence regions contributing to predictions at individual binding peaks, termed *seqlets*, and clusters these into recurrent patterns summarized as contribution weight matrices (CWMs). These patterns often correspond to individual TF-binding motifs, as well as composite motifs representing combinations of binding sites that could reflect cooperative TF interactions (Fig. 1E). As a first test of motif cooperativity, we examined the prevalence of composite patterns across TF models^7^. Although detected, these patterns generally occurred in only a minority of binding peaks. Moreover, individual seqlets typically contained limited sequence information, approximately 12 bits, equivalent to a fully specified 6-bp sequence^12^ (Fig. 1 E-I). Together, these findings suggest that neither composite motifs nor high-information-content sequence patterns contribute substantially to model-predicted binding specificity at the genomic scale.

Given the limited information content of individual seqlets, we next asked whether binding specificity emerges from cooperative interactions among multiple motifs within the same binding peak. We first focused on multiple occurrences of the motif bound by the profiled TF (direct motif), reasoning that occupancy at one site may enhance binding at neighboring sites^7,13^. To test this, we performed *in silico* perturbations, comparing the effects of mutating motifs individually or in combination (Fig. 2A-C). Under independent contributions, the effect of the combined mutations is expected to equal the sum of the individual effects, whereas a larger or smaller summed effect indicates cooperativity or redundancy, respectively^14,15^. Note that we used control-free predicted count scores, as BPNet predictions were often dominated by the free-DNA control (Fig. 1D).

**Figure 2.**
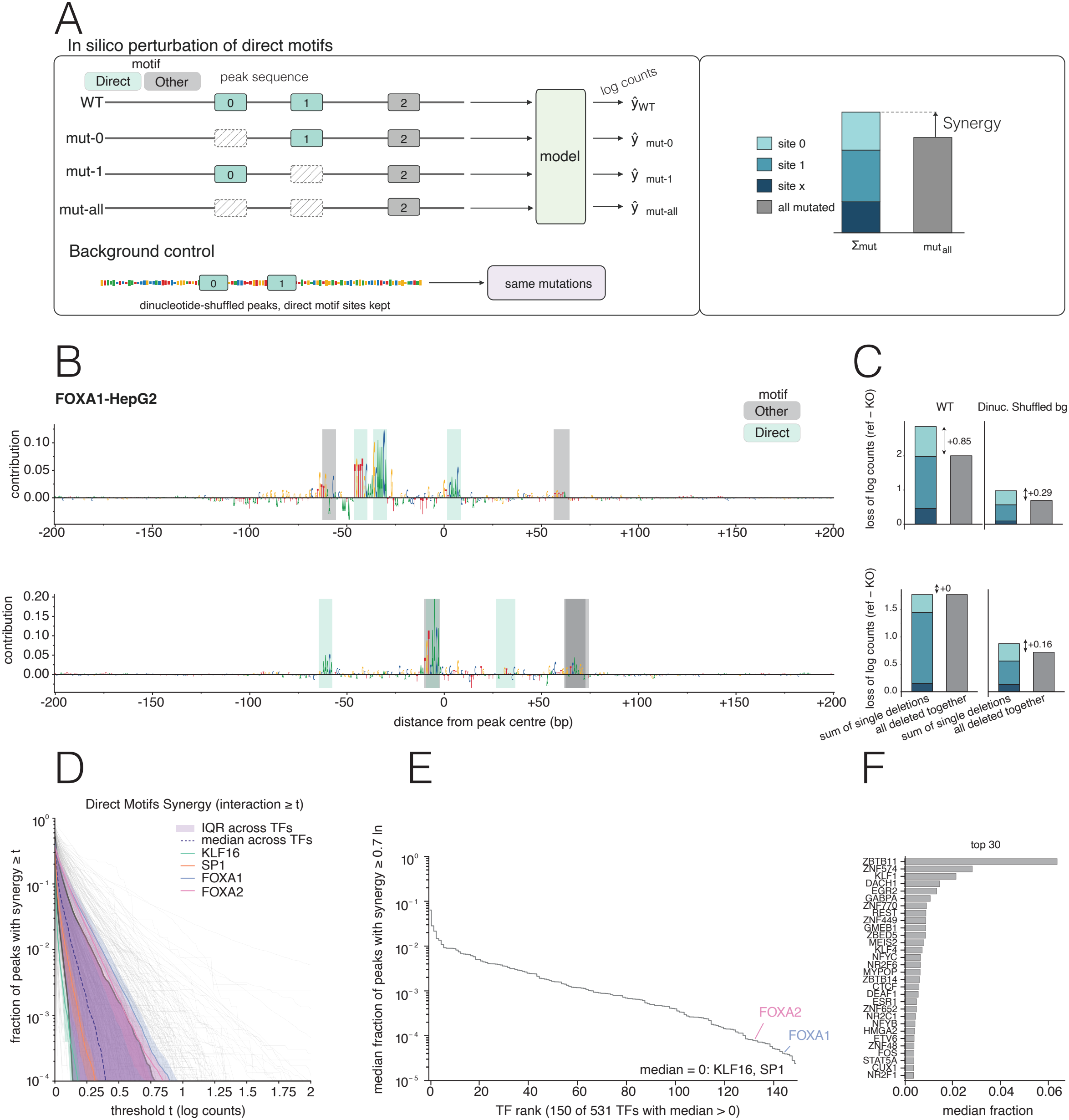
Binding peaks often contain multiple seqlets corresponding to self-motifs, but their contributions are predominantly additive or partially redundant, while cooperativity is rare. **(A)** Schematic of the *in silico* perturbation strategy used to assess interactions between seqlets corresponding to direct (self) motifs within individual binding peaks. Predicted count scores were obtained for the wild-type (WT) sequence, sequences with each direct motif site individually mutated (mut-0 and mut-1), and a sequence in which all direct motif sites were mutated (mut-all). Synergy was assessed by comparing the summed effects of individual motif mutations with the effect of the combined mutation. Under independent motif contributions, the combined effect is expected to equal the sum of the individual effects. A larger summed individual effect indicates cooperativity, whereas a smaller summed effect indicates redundancy. As a background control, dinucleotide-shuffled peak sequences retaining the direct motif sites were subjected to the same perturbations. Throughout the analysis, we used log-count scores to quantify binding and define additive and non-additive motif contributions. Notably, cooperative effects remained limited when evaluated in linear-count space (Fig. S3), despite its greater tendency to classify multiplicative motif contributions as cooperative, because motif contributions were often redundant or had only small effects on predicted binding. **(B)** Contribution profiles around two representative FOXA1-HepG2 binding peaks. Direct motifs and other motifs identified by the model are indicated, together with their contribution scores and positions relative to the peak center. **(C)** Example comparisons of the effects of individual versus combined direct motif mutations on predicted binding, shown for WT peak sequences and corresponding dinucleotide-shuffled background controls. Bars represent the loss of predicted log counts relative to the reference sequence, with contributions from individual motif mutations and their combined mutation shown separately. **(D)** Fraction of binding peaks exhibiting cooperative interactions between direct motifs above increasingly stringent synergy thresholds, shown across TF-averaged models. The median and interquartile range (IQR) across TFs are indicated. Selected TFs, including KLF16, SP1, FOXA1, and FOXA2, are highlighted, with the standard error of the mean across all experiments for the respective TF as a ribbon. **(E)** TF models ranked according to the median fraction of peaks exhibiting synergy above two-fold. The analysis includes 150 of 531 TFs with a median fraction greater than zero. FOXA1 and FOXA2 are highlighted. **(F)** The 30 TF models with the highest median fractions of peaks exhibiting cooperative interactions between direct motifs above the indicated threshold.

Overall, cooperative interactions among direct motifs made a negligible contribution to model-predicted TF binding (Fig. 2D-F). Across peaks and TFs, motif contributions were largely additive. Cooperative effects were detected at individual peaks and were more prominent for certain TFs, including FOXA1 and FOXA2. However, their prevalence decreased sharply with increasing effect size, with effects exceeding twofold observed in fewer than 1% of peaks for nearly all TFs (Fig. 2D-F).

We next asked whether interactions between motifs of the profiled TF and those of other TFs (non-direct motifs) might play a more prominent role in predicted binding. To investigate this, we extended our *in silico* perturbation analysis to direct and non-direct motifs (Fig. 3A-C), comparing the effects of mutating either group individually or both together at each binding peak. Notably, some TFs showed greater dependence on non-direct motifs, although direct motifs dominated in most cases (Fig. 3F-G). We then assessed whether the contribution of one motif group depended on the presence of the other, as expected under cooperative binding. As with interactions among direct motifs, the combined effects were predominantly additive, with cooperative interactions detected only in a minority of cases (Fig. 3D-E). Thus, cooperativity between direct and non-direct motifs made a negligible contribution to genome-wide binding predictions.

**Figure 3.**
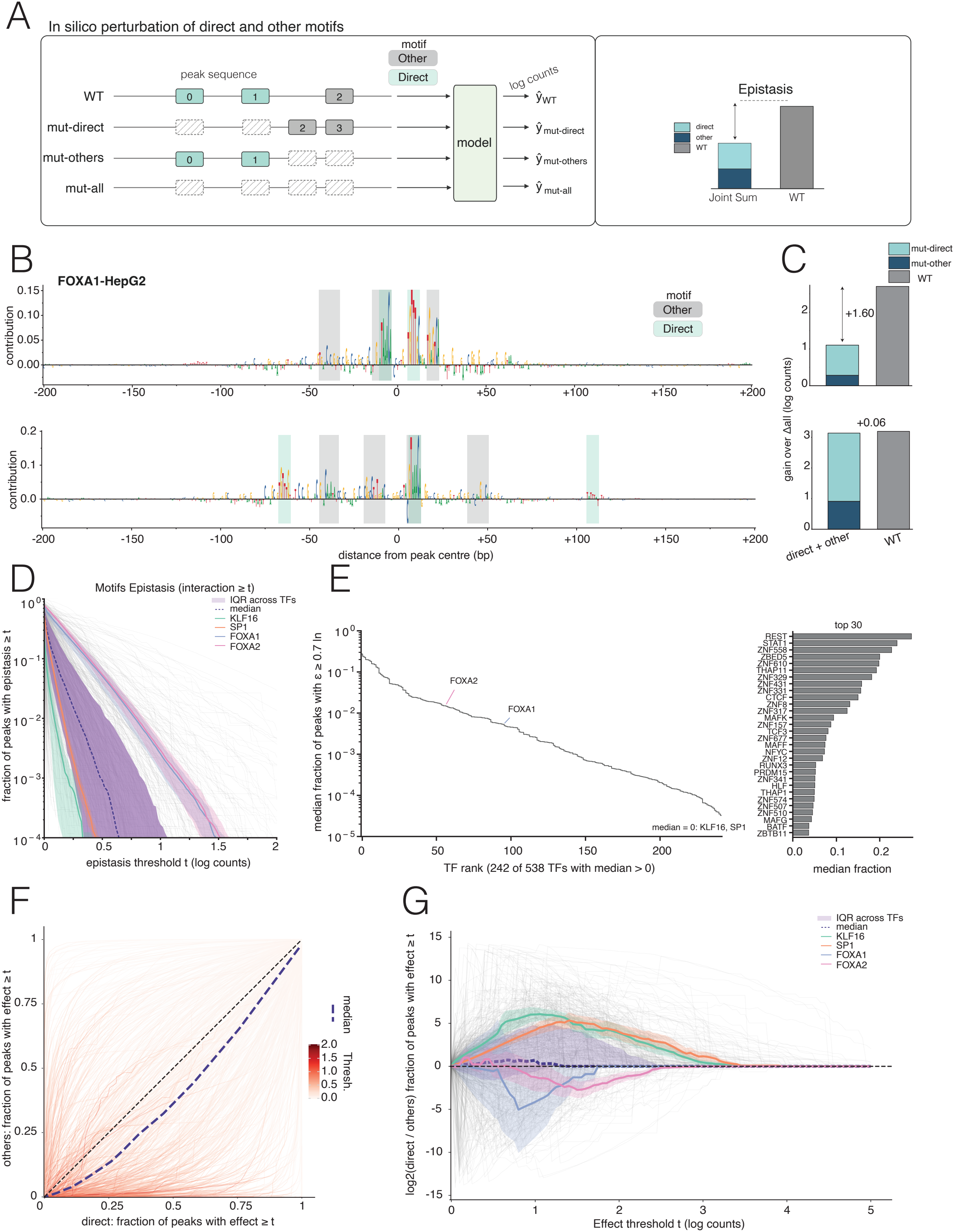
Binding peaks often contain multiple seqlets corresponding to direct and non-self motifs, but their contributions are predominantly additive or partially redundant, while epistatic interactions are rare. **(A)** Schematic of the in silico perturbation strategy used to assess interactions between direct (self) and other (non-self) motifs within individual binding peaks. Predicted log counts are obtained for the WT sequence, sequences with direct motifs mutated (mut-direct), sequences with other motifs mutated (mutothers), and sequences in which both motif groups are mutated (mut-all). Epistasis is assessed by comparing the combined effect of mutating both motif groups with the joint contribution of the individual mutations. Deviations from the expected additive effect indicate dependencies between direct and other motifs. **(B)** Contribution profiles around two representative FOXA1-HepG2 binding peaks. Direct and other motifs identified by the model are indicated, together with their contribution scores and positions relative to the peak center. **(C)** Representative comparisons of the effects of deleting direct motifs, other motifs, or both groups together on predicted binding. The gain in predicted log counts following the joint mutation, relative to the combined mutation condition (mut-all), is shown for each example. **(D)** Fraction of binding peaks exhibiting epistatic interactions between direct and other motifs above increasingly stringent thresholds, shown across TF-averaged models (presentation as in Fig. 2D). **(E)** TF models ranked according to the median fraction of peaks exhibiting epistasis above 0.7 log counts. The analysis includes 242 of 538 TFs with a median fraction greater than zero. FOXA1 and FOXA2 are highlighted. **(F)** Relationship between the fraction of peaks showing effects above a given threshold following mutation of direct motifs and the corresponding fraction following mutation of other motifs. Each point represents a TF model, with color indicating the effect threshold. The dashed diagonal denotes equal contributions from the two motif groups, and the purple diagonal the median effect of all TFs. **(G)** Difference in the prevalence of direct versus other motif contributions across increasingly stringent effect thresholds, expressed as the log□ ratio of the fractions of peaks affected by mutation of direct motifs and other motifs. The median and IQR across TFs are indicated, together with selected TFs.

Next, we asked how TFs with the same motif preferences bind in the same cells and chromatin context, and whether the models capture that. If binding preferences were determined primarily by canonical motif recognition and chromatin accessibility, TFs recognizing similar motifs within the same cellular environment would be expected to occupy similar genomic sites. To examine this, we systematically compared genome-wide binding profiles of TFs belonging to the same motif family and profiled in the same cell type across the ENCODE dataset. Notably, as exemplified by the FOX and SP/KLF families (Fig. 4A-C), TFs sharing similar motif preferences often exhibited substantially different genomic binding profiles (Fig. S2). These differences were accurately captured by the sequence-based models and persisted when mutating the major seqlets (Fig. 4D-E). Thus, canonical motif recognition and a shared chromatin environment are insufficient to fully explain TF genomic selectivity, pointing to additional sequence determinants captured by the models.

**Figure 4.**
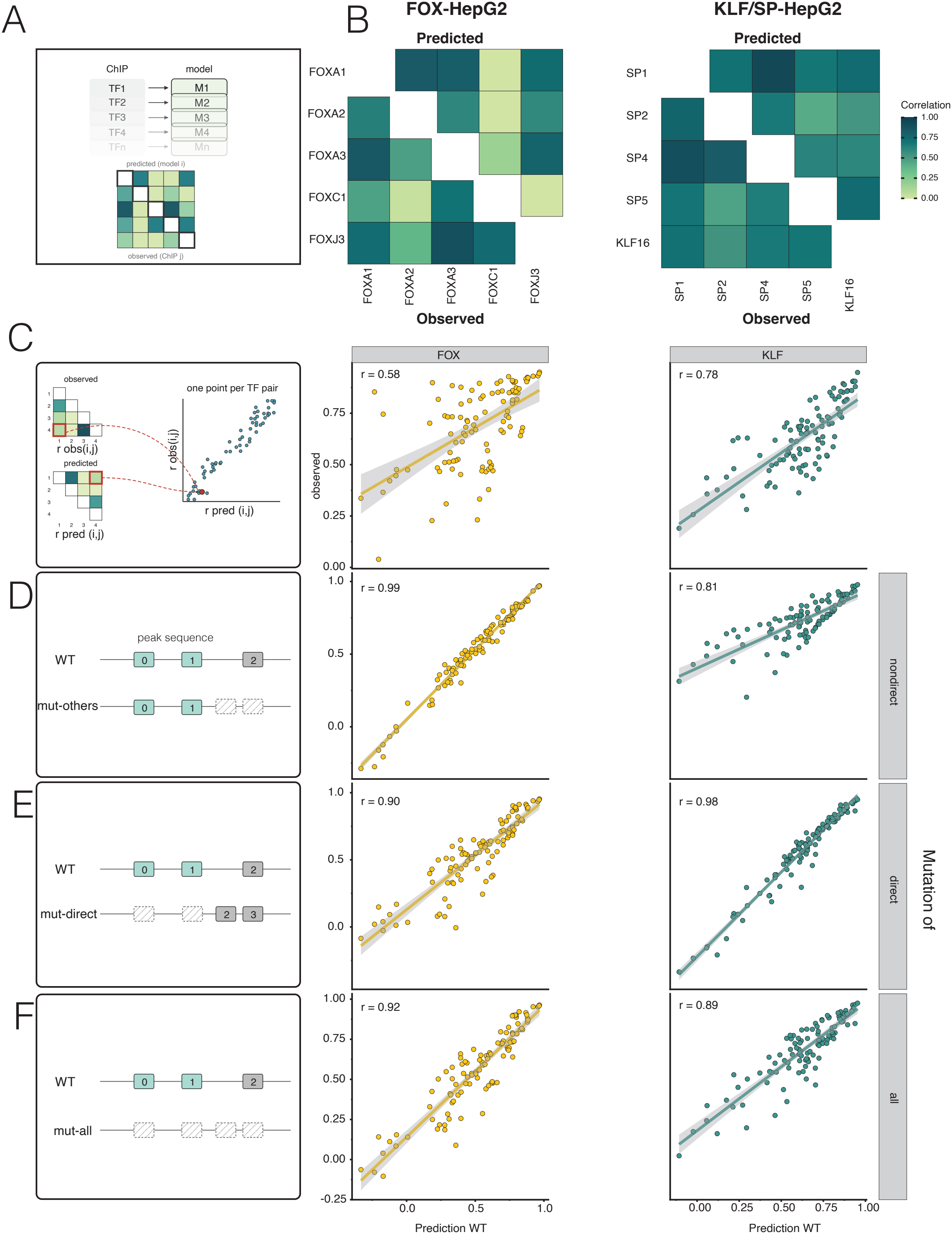
Sequence-based models distinguish binding preferences of TFs with similar motif recognition within the same cellular environment. **(A)** Schematic of the cross-prediction framework used to compare TF binding preferences. Each BPNet model, trained on ChIP-seq data for a particular TF, is used to predict binding for genomic sequences profiled for multiple TFs. The similarity between binding profiles of the predicted and observed binding profiles is quantified using Pearson correlation and is shown for the observed binding (lower triangle) and the predicted binding (upper triangle), allowing comparisons between observed and predicted TF similarities. **(B)** Heatmaps comparing predicted and observed binding profiles for TFs belonging to the FOX family (left, 5/15 experiments in HepG2 cell line) and the KLF/SP family (right, 5/15 experiments in HepG2 cell line), profiled in HepG2 cells, organized as in A. **(C)** Relationship between pairwise similarities in observed TF binding profiles and pairwise similarities in model-predicted profiles for the FOX family (left, all 15 experiments presented) and KLF/SP family (right, all 15 experiments presented). Each point represents a pair of TFs. Correlations between observed and predicted pairwise similarities are indicated. **(D–F)** Comparison of WT and in silico-perturbed model predictions for the FOX family (left, all 15 experiments presented) and KLF/SP family (right, all 15 experiments presented). Schematic representations of the sequence perturbations are shown on the left of each panel. Each point represents a binding peak, with predicted WT binding on the horizontal axis and predicted binding following sequence perturbation on the vertical axis. Pearson correlations between WT and perturbed predictions are indicated. (D) Mutation of other (non-direct) motifs. (E) Mutation of direct motifs. (F) Mutation of all identified motifs.

Together, our analysis reveals that sequence-based models accurately predict selective TF binding with limited reliance on motif cooperativity and that chromatin accessibility alone is insufficient to explain genomic preferences. Importantly, accurate prediction of TF binding profiles from DNA sequence does not necessarily imply that models capture the underlying biological mechanisms, as their learned representations may reflect alternative solutions, biases in ChIP-seq data, or other experimental confounders. Nevertheless, our findings, together with other studies questioning the central role of cooperativity in genomic targeting^15-17^ raise the possibility that additional specificity mechanisms remain uncharacterized. For example, specificity may depend on weak, distributed DNA sequence signals extending beyond recognizable motifs^18,19^, potentially paralleling the distributed binding determinants identified within TF intrinsically disordered regions (IDRs), which are increasingly recognized as important contributors to genomic binding preference^20-24^. Distinguishing among these possibilities will require further experimental and computational investigation to establish the molecular basis of selective TF binding.

## Acknowledgements

We thank our past and current lab members for technical assistance, fruitful discussions, and comments on the manuscript. Vladimir Mindel is supported by the Adams Fellowships of the Israel Academy of Sciences and Humanities.

## Code and data availability

The code and notebooks needed to reproduce the analysis are available under the following accession: DOI: 10.5281/zenodo.24280380

