## Supplementary Figures and Methods for "Sequence-based models predict TF binding with negligible contributions from motif cooperativity"

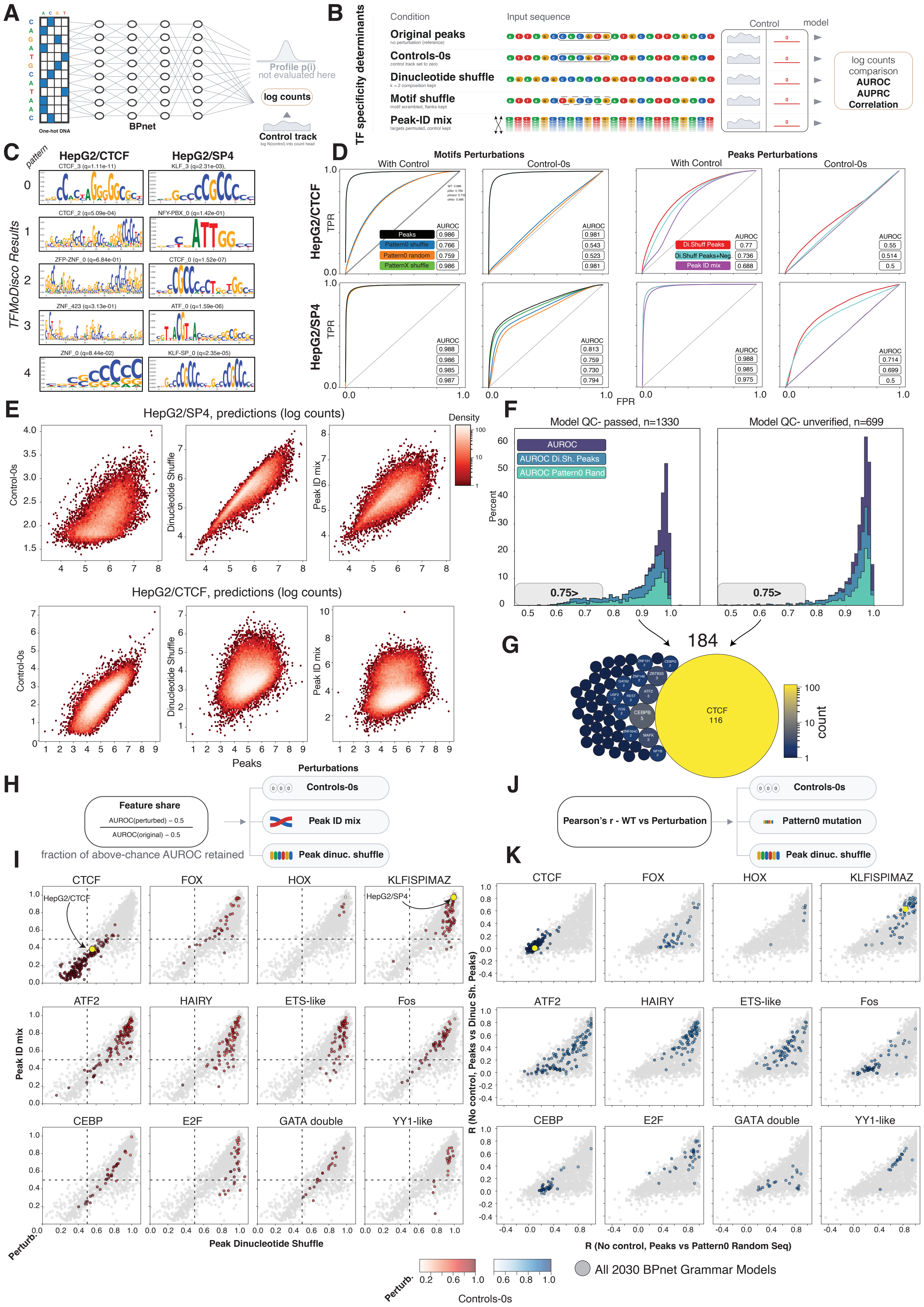
A
One-hot DNA
BPnet
Profile p(i)
not evaluated here
log counts
Control track
B
TF specificity determinants
Condition
Input sequence
Control
model
Original peaks
no perturbation (reference)
Controls-0s
control track set to zero
Dinucleotide shuffle
k = 2 composition kept
Motif shuffle
motif scrambled, flanks kept
Peak-ID mix
targets permuted, control kept
log counts comparison
AUROC
AUPRC
Correlation
C
HepG2/CTCF
HepG2/SP4
pattern
TFMoDisco Results
CTCF_3 (q=1.11e-11)
KLF_3 (q=2.31e-03),
CTCF_2 (q=5.09e-04)
NFY-PBX_0 (q=1.42e-01)
ZFP-ZNF_0 (q=6.84e-01)
CTCF_0 (q=1.52e-07)
ZNF_423 (q=3.13e-01)
ATF_0 (q=1.59e-06)
ZNF_0 (q=8.44e-02)
KLF-SP_0 (q=2.35e-05)
D
Motifs Perturbations
Peaks Perturbations
With Control
Control-0s
HepG2/CTCF
HepG2/SP4
TPR
FPR
Peaks
Pattern0 shuffle
Pattern0 random
PatternX shuffle
Di.Shuff Peaks
Di.Shuff Peaks+Neg.
Peak ID mix
AUROC
E
HepG2/SP4, predictions (log counts)
HepG2/CTCF, predictions (log counts)
Control-0s
Dinucleotide Shuffle
Peak ID mix
Peaks
Density
F
Model QC- passed, n=1330
Model QC- unverified, n=699
AUROC
AUROC Di.Sh. Peaks
AUROC Pattern0 Rand
Percent
0.75>
G
184
CTCF
116
count
H
Feature share
AUROC(perturbed) − 0.5
AUROC(original) − 0.5
fraction of above-chance AUROC retained
Perturbations
Controls-0s
Peak ID mix
Peak dinuc. shuffle
J
Pearson's r - WT vs Perturbation
Controls-0s
Pattern0 mutation
Peak dinuc. shuffle
I
CTCF
FOX
HOX
KLFISPIMAZ
HepG2/CTCF
HepG2/SP4
ATF2
HAIRY
ETS-like
Fos
CEBP
E2F
GATA double
YY1-like
Peak ID mix
Peak Dinucleotide Shuffle
Perturb.
K
R (No control, Peaks vs Dinuc Sh. Peaks)
R (No control, Peaks vs Pattern0 Random Seq)
All 2030 BPnet Grammar Models
Controls-0s

**Figure S1. Assessing the sequence and control determinants of BPNet binding predictions across ENCODE TF models.**

**(A) Schematic representation of the BPNet architecture:** The model receives one-hot encoded DNA sequences and experimental control tracks as inputs and predicts base-resolution binding profiles and total binding strength (log counts). Only the predicted log-count output was evaluated in our analysis.

**(B) Schematic of the *in silico* perturbations used to assess the contribution of sequence and control information:** The following conditions were examined: original peak sequences provided with their corresponding control tracks (reference), original sequences with control tracks set to zero (Controls-0s), dinucleotide shuffling of the input DNA sequence (preserving dinucleotide composition), motif shuffling (scrambling the identified motif while preserving flanking sequences), and peak-ID mixing (permuting target peak identities while retaining the control track). Model predictions were evaluated by comparing predicted and observed binding using log-counts, AUROC, AUPRC, and correlation.

**(C) Representative TF-MoDISco contribution weight matrices (CWMs):** Shown are CWMs identified in the HepG2 CTCF and SP4 models, illustrating sequence patterns contributing to model predictions.

**(D) Sensitivity of model predictions to input perturbations, quantified by receiver operating characteristic (ROC) curves:** Shown are ROC curves comparing the effects of motif perturbations and peak-level sequence perturbations on binding classification in HepG2 CTCF and SP4 models, with and without control tracks. AUROC values are indicated.

**(E) Sensitivity of predicted binding strength to input perturbations, quantified by count-score correlations:** Scatter plots compare predicted log-counts for original peaks (x-axis) with predictions following control removal, dinucleotide shuffling, or peak-ID mixing (y-axis). Results are shown for HepG2 SP4 (top) and CTCF (bottom). Each point represents a genomic region, and color indicates point density.

**(F-G) Quantifying the sensitivity of the model's prediction to input perturbations across the ENCODE BPNet model collection:** Shown in (G) are distributions of AUROC values across BPNet models with passed quality control (n = 1,330) and unverified quality control status (n = 699), comparing original predictions with predictions following dinucleotide shuffling of peak sequences and randomization of the primary sequence pattern (Pattern 0). The dashed line indicates AUROC = 0.75; the AUROC across TFs is also shown (G), with bubble size and color representing the number of models associated with each TF.

**(H–I) Sensitivity of model predictions to control and DNA sequence information differs across TF families:** We measured the fraction of above-chance AUROC retained following the three indicated input perturbations, as illustrated schematically (H). Results are shown across ENCODE BPNet models grouped by TF motif family, comparing the effects of peak-ID mixing and dinucleotide shuffling, with colors indicating the effect of control removal (I). Higher values indicate lower sensitivity to the corresponding perturbation.

**(J–K) Sensitivity of model predictions to DNA sequence perturbations in the absence of control input differs across TF families:** We measured the Pearson correlation between predicted log-counts for original and perturbed sequences, with control tracks set to zero, as illustrated schematically (J). Results are shown across ENCODE BPNet models grouped by TF motif family, comparing the effects of primary motif (Pattern 0) randomization and dinucleotide shuffling of the entire peak sequence (K). Higher correlations indicate lower sensitivity to the corresponding sequence perturbation.

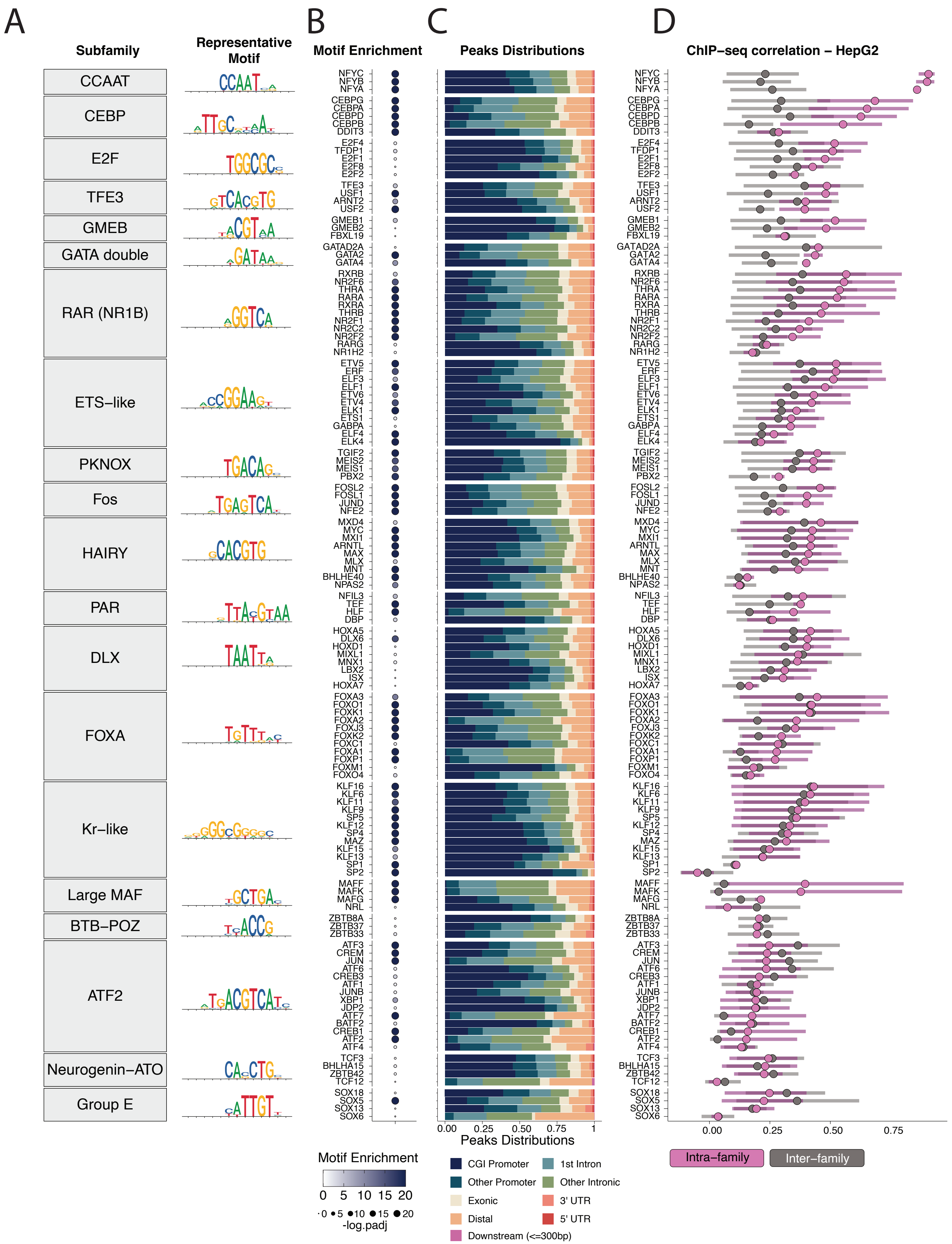
A
Subfamily
Representative Motif
CCAAT
CEBP
E2F
TFE3
GMEB
GATA double
RAR (NR1B)
ETS–like
PKNOX
Fos
HAIRY
PAR
DLX
FOXA
Kr–like
Large MAF
BTB–POZ
ATF2
Neurogenin–ATO
Group E
B
Motif Enrichment
C
Peaks Distributions
D
ChIP–seq correlation – HepG2
NFYC
NFYB
NFYA
CEBPG
CEBPA
CEBPD
CEBPB
DDIT3
E2F4
TFDP1
E2F1
E2F8
E2F2
TFE3
USF1
ARNT2
USF2
GMEB1
GMEB2
FBXL19
GATAD2A
GATA2
GATA4
RXRB
NR2F6
THRA
RARA
RXRA
THRB
NR2F1
NR2C2
NR2F2
RARG
NR1H2
ETV5
ERF
ELF3
ELF1
ETV6
ETV4
ELK1
ETS1
GABPA
ELF4
ELK4
TGIF2
MEIS2
MEIS1
PBX2
FOSL2
FOSL1
JUND
NFE2
MXD4
MYC
MXI1
ARNTL
MAX
MLX
MNT
BHLHE40
NPAS2
NFIL3
TEF
HLF
DBP
HOXA5
DLX6
HOXD1
MIXL1
MNX1
LBX2
ISX
HOXA7
FOXA3
FOXO1
FOXK1
FOXA2
FOXJ3
FOXK2
FOXC1
FOXA1
FOXP1
FOXM1
FOXO4
KLF16
KLF6
KLF11
KLF9
SP5
KLF12
SP4
MAZ
KLF15
KLF13
SP1
SP2
MAFF
MAFK
MAFG
NRL
ZBTB8A
ZBTB37
ZBTB33
ATF3
CREM
JUN
ATF6
CREB3
ATF1
JUNB
XBP1
JDP2
ATF7
BATF2
CREB1
ATF2
ATF4
TCF3
BHLHA15
ZBTB42
TCF12
SOX18
SOX5
SOX13
SOX6
0.00 0.25 0.50 0.75 1
Peaks Distributions
0.00 0.25 0.50 0.75
Intra–family
Inter–family
Motif Enrichment
0 5 10 15 20
0 5 10 15 20
-log.padj
CGI Promoter
1st Intron
Other Promoter
Other Intronic
Exonic
3' UTR
Distal
5' UTR
Downstream (<=300bp)

**Figure S2. Systematic comparison of genomic binding profiles among TFs sharing similar DNA-binding motifs.**

**(A) Motif-sharing TFs profiled:** We classified ENCODE-profiled TFs into motif-sharing families based on their canonical recognition motifs, following the HOCOMOCOv14 database classification. All families with at least two members profiled in HepG2 cells are shown, together with their respective consensus motifs. All analyses in this figure were performed using ENCODE HepG2 data, with similar results observed in other cell types.

**(B) Enrichment of consensus motifs within TF-binding peaks:** Enrichment of the corresponding consensus motif within the binding peaks of each TF is quantified by the −log10 adjusted p-value and represented by dot size.

**(C) Genomic distribution of TF-binding peaks:** The distribution of ChIP-seq peaks across annotated genomic regions is shown for each TF, with colors indicating the fraction of peaks found in CpG island promoters, other promoters, exons, distal regions, introns, untranslated regions (UTRs), and other annotated regions.

**(D) Motif-sharing TFs often exhibit limited similarity in genomic binding:** We measured the correlation between each TF's genomic binding profile and those of other TFs sharing its recognition motif (intra-family), as well as all other profiled TFs (inter-family). The respective distributions of correlation values are compared for each TF. While components of the same TF complex exhibit high binding correlations (e.g., NFY complex), members of larger motif-sharing families, including FOX and KLF, generally do not exhibit greater intra-family similarity than their correlations with TFs outside their respective families.

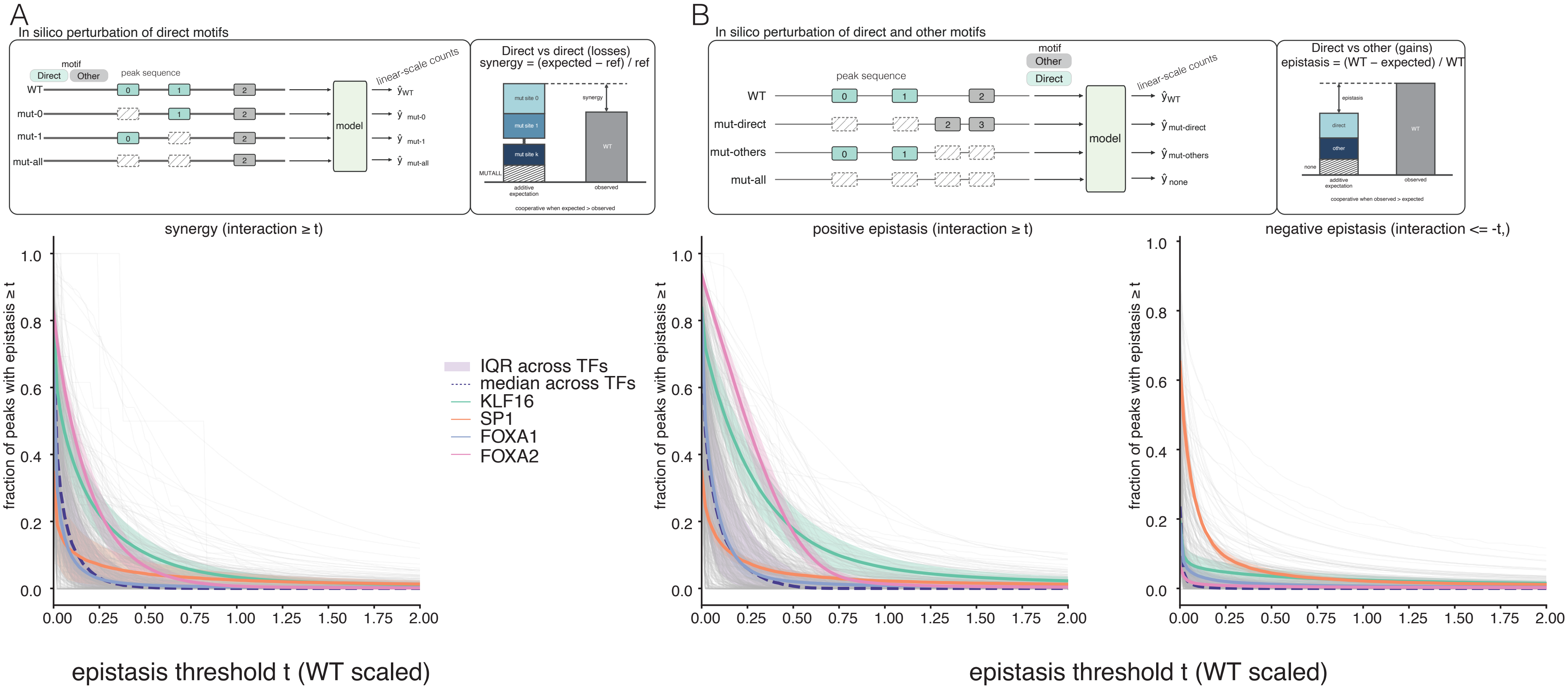
A
In silico perturbation of direct motifs
motif
Direct
Other
peak sequence
linear-scale counts
WT
mut-0
mut-1
mut-all
model
Direct vs direct (losses)
synergy = (expected – ref) / ref
mut site 0
mut site 1
mut site k
synergy
WT
MUTALL
additive expectation
observed
cooperative when expected > observed
B
In silico perturbation of direct and other motifs
mut-direct
mut-others
Direct vs other (gains)
epistasis = (WT – expected) / WT
epistasis
direct
other
none
cooperative when observed > expected
synergy (interaction ≥ t)
positive epistasis (interaction ≥ t)
negative epistasis (interaction <= -t,)
fraction of peaks with epistasis ≥ t
IQR across TFs
median across TFs
KLF16
SP1
FOXA1
FOXA2
epistasis threshold t (WT scaled)

**Figure S3. Binding peaks often contain multiple seqlets corresponding to direct and non-self motifs, but their contributions are predominantly additive or partially redundant on a linear scale.**

**(A)** Schematic of the *in silico* perturbation strategy used to assess interactions between seqlets corresponding to direct (self) motifs within individual binding peaks**,** in linear scale (top, presentation as in Fig. 2A), and the fraction of binding peaks exhibiting cooperative interactions between direct motifs above increasingly stringent synergy thresholds, shown across TF-averaged models (bottom, presentation as in Fig. 2D). Note: Exceptionally high apparent cooperativity scores exceeding 1 arise only when the expected ordering of predicted binding signals (WT > mut-A, mut-B > mut-AB) is violated. Such violations may also occur at elevated scores below 1, but this was not systematically assessed.

**(B)** Schematic of the *in silico* perturbation strategy used to assess interactions between seqlets corresponding to direct (self) and other (non-self) motifs within individual binding peaks**,** in linear scale (top, presentation as in Fig. 3A), and the fraction of binding peaks exhibiting cooperative interactions between self and non-self motifs above increasingly stringent synergy thresholds, shown across TF-averaged models (bottom, presentation as in Fig. 3D)

### Methods

#### BPNet models and motif annotations

BPNet models (Avsec et al. 2021) trained on ENCODE TF ChIP-seq experiments (ENCODE Project Consortium 2026; Yun et al. 2026) were obtained through the ENCODE API. The released model was used where available; for the remaining experiments, we used the archived model, i.e., one that did not pass ENCODE quality control, and these models were flagged. Each model comprises five networks, each trained with a different set of held-out chromosomes. For each experiment, we downloaded the models, the training and test regions, the conservative and pooled-pseudoreplicate IDR peaks, the observed control-track bigWig files, the TF-MoDISco motif reports and matrices, the Fi-NeMo motif instances, and the ENCODE-annotated motif-instance bigBed files. The bigBed files classify each motif instance relative to the profiled TF as direct, indirect, non-canonical, or unvalidated. Throughout, “direct motifs” refers to instances of positive counts-head patterns annotated as direct, and “other motifs” to all remaining positive counts-head instances. Unless stated otherwise, predictions are the counts-head output (predicted log counts), obtained with the control track set to zero and averaged over the five folds.

#### Model performance with and without the control track

To determine how much of the performance of each model depends on the input control track (Fig. 1D, Fig. S1), we reconstructed the ENCODE test sets for each model. The test peaks of each fold were 2,114-bp windows centered on the peak summit, and GC-matched background regions were taken from the ENCODE training and test region files. As in the ENCODE procedure, background regions overlapping the IDR-ranked peaks of the experiment were removed. Regions were scored by the network of the fold in which they were held out, once using the observed control profile of each region and once using an all-zero control track. Performance was quantified as the area under the receiver operating characteristic curve (AUROC) for separating peaks from background regions by predicted log counts. The AUROC values obtained with the control track reproduced those reported by ENCODE. Values were averaged over the five folds, and their distributions were compared across all 2,282 models.

#### Information content of motif cores

Differences in information content and frequency between motifs identified by the same model are illustrated for the BHLHE40 model in K562 cells (ENCSR000EGV; Fig. 1). Sequence logos show the information content of two of its untrimmed TF-MoDISco counts-head motifs, pattern_0 and pattern_8, and the total information content of each matrix was summed over all positions. The fraction of peaks containing each motif was calculated using 1-kb windows centered on the peak summits; a window was considered to contain a motif if it overlapped at least one Fi-NeMo instance of that motif ($\lambda = 0.7$).

For all models, the information content of each untrimmed TF-MoDISco counts-head motif was calculated over its central 20 bp (the motif core). Most ENCODE motifs are 50 bp wide, corresponding to a 20-bp TF-MoDISco seqlet flanked by 15 bp on each side. 58 models, mostly of zinc-finger TFs, have 110-bp motifs; for these, the central 20 bp of the 30-bp seqlet were used, so that all cores were 20 bp long. Core information content was expressed as effective length in base pairs (bp), i.e., bits divided by two. For each motif, we also recorded the fraction of the peaks of the model containing at least one Fi-NeMo instance (counts head, $\lambda = 0.7$). Motifs were grouped by TF-MoDISco rank (patterns 1–5, 6–10 and beyond 10). Each model was summarized by the mean effective core length of its motifs, weighted by the number of peaks in which each motif occurred, and models of the same TF were averaged. This per-TF value was compared with the core information content of the top-ranked motif (pattern_0) of each TF using Pearson and Spearman correlations ($n$ = 868 TFs).

**In silico knockout of direct and other motifs**

The contributions of direct and other motifs to predicted binding, alone and in combination, were measured by in silico knockouts in which motif bases were replaced by shuffled donor sequence (Figs. 2, 3; Fig. S3). Each peak of each model was centered within a 2,114-bp window, and windows containing N were excluded. Motif instances from the ENCODE bigBed files that overlapped the central 600 bp were assigned to the direct or the other group; bases covered by both groups were assigned to the direct group. For each peak, 20 dinucleotide-shuffled versions of the central 600 bp served as donors. In a knockout, the bases of one motif group were replaced by the corresponding bases of a donor, and all remaining bases were left unchanged. Predictions were obtained for four versions of each peak: the intact sequence ($P_{\mathrm{wt}}$), the sequence with both groups knocked out ($P_{\mathrm{none}}$), the sequence retaining only the direct motifs ($P_{\mathrm{direct}}$; mut-others in the figures) and the sequence retaining only the other motifs ($P_{\mathrm{other}}$; mut-direct in the figures). Predictions for knocked-out sequences were averaged over the 20 donors. Epistasis between the two motif groups was defined as

$$I = (P_{\mathrm{wt}} - P_{\mathrm{none}}) - [(P_{\mathrm{direct}} - P_{\mathrm{none}}) + (P_{\mathrm{other}} - P_{\mathrm{none}})],$$

where $P$ denotes predicted log counts. $I$ is thus the difference between the effect of both groups in the intact peak and the additive expectation from each group alone.

**Knockout of individual direct sites**

To test whether the direct sites within a peak act independently (Fig. 2), each direct site was knocked out individually and all direct sites were knocked out together. Donors were the same as above (20 dinucleotide shuffles of the central 600 bp, generated with the same random seeds). Direct sites sharing at least one base were first merged into a single site spanning their union. Knockouts were performed in two sequence contexts: the native peak (WT), in which each site was replaced by donor sequence in the otherwise intact peak, and a shuffled background, in which the central 600 bp were shuffled except for the direct sites and each site was then also removed from this background. For peaks with at least two direct sites, the synergy among them was defined as

$$S = \sum_{i} \left(P_{\mathrm{ref}} - P_{\mathrm{KO},i}\right) - \left(P_{\mathrm{ref}} - P_{\mathrm{KO,all}}\right),$$

where $P_{\mathrm{ref}}$ is the prediction in the given context, $P_{\mathrm{KO},i}$ the prediction with site $i$ removed, and $P_{\mathrm{KO,all}}$ the prediction with all direct sites removed. $S$ therefore corresponds to the sum of the single-site losses minus the joint loss.

**Epistasis and synergy across TFs**

For each experiment, we calculated the fraction of peaks in which the interaction $I$ (Fig. 3) or the direct-site synergy $S$ in the native context (Fig. 2) was at least $t$, for thresholds $t$ from 0 to 2 log counts in steps of 0.02. The interaction was evaluated across peaks containing both direct and other motifs (1,480 experiments, 538 TFs), and synergy was evaluated across peaks containing at least two direct sites (1,506 experiments, 531 TFs). To prevent TFs profiled in many experiments from dominating, curves were averaged per TF. The curves of all TFs are shown together with the median and interquartile range (IQR) across TFs, and with the mean ± SEM for selected TFs. TFs were ranked by the median, across their experiments, of the fraction of peaks exceeding $t = 0.7$ log counts.

**Individual effects of direct and other motifs**

Over the same peaks, we calculated for each experiment the fraction of peaks in which the direct motifs alone ($f_{\mathrm{direct}}$) or the other motifs alone ($f_{\mathrm{other}}$) increased the prediction by at least $t$, for $t$ from 0 to 5 log counts in steps of 0.05 (Fig. 3). Per-TF means of the two fractions were plotted against each other across thresholds. Their balance was summarized as

$$R = \log_2 \frac{f_{\text{direct}} + \varepsilon}{f_{\text{other}} + \varepsilon}, \qquad \varepsilon = 5 \times 10^{-5}.$$

$R$ was calculated per experiment and then averaged per TF, and is shown for $t \leq 5$ together with the median and IQR across TFs.

**Binding similarity among KLF/SP and FOX family members**

To examine why members of a TF family bind differently (Fig. 4), we analyzed 15 KLF/SP and 15 FOX BPNet models trained on HepG2 experiments. For each family, the conservative IDR peaks of all members were merged into one set, and each merged region was centered in a 2,114-bp window; windows containing N or extending beyond the chromosome end were excluded. For each member, the observed signal was summed over the central 1 kb of each window from the plus- and minus-strand observed signal bigWig files and log1p-transformed, matching the target of the counts head. The predicted signal was the predicted log counts of each member over the same windows, with the control track set to zero. Motif instances were assigned to the family motif if they were annotated as direct by ENCODE or matched a family position weight matrix (PWM) at $P < 10^{-4}$. PWMs were taken from HOCOMOCO v14 (Vorontsov et al. 2024): SP1 and KLF16 for KLF/SP, and seven FOX PWMs for FOX. Within the central 1 kb, we knocked out the family-motif bases, the other-motif bases or all motif bases; each knockout used five dinucleotide-shuffled donors and was predicted by the models of all members.

Similarity between members was defined as the Pearson correlation of their signals across windows and was calculated for each pair of members (105 pairs per family) from the observed signal, the predicted signal and the predictions after each knockout. Correlation matrices show the observed and predicted similarity for five representative members, with the correlation between the observed and predicted signal of each member on the diagonal. Scatter plots compare the similarity of each pair after each knockout with its similarity in the intact prediction.

**Contribution of the control track, motifs and sequence composition to model performance**

To separate the contributions of the control track, the motifs and sequence composition to model performance (Fig. S1), the ENCODE test folds described above were scored under a series of sequence perturbations. Each perturbation was evaluated twice, with the observed control profile of each region and with an all-zero control track, and each region was scored only by the network of the fold in which it was held out. First, within each fold, every region (peaks and background together) was assigned the sequence of another region, selected by a random derangement (seed 0), while retaining its own control profile (identity mix). Second, the full 2,114-bp sequence of each region was dinucleotide-shuffled once, which destroys motifs while preserving dinucleotide and GC composition; shuffled peaks were compared with the unshuffled background and with shuffled background. Third, motifs were knocked out in the test peaks using the ENCODE Fi-NeMo instances (counts head, $\lambda = 0.7$) that overlapped the 2,114-bp window of each peak, while background regions were left unchanged. The bases of all pattern_0 instances (the top-ranked TF-MoDISco motif) were either permuted, which preserves the base composition of the window (p0ko), or replaced by uniformly random bases, which also changes the composition (p0rand). In addition, the bases of all other positive-pattern instances were permuted while the pattern_0 positions were kept fixed (othko).

For each perturbation, the AUROC and the area under the precision–recall curve (AUPRC) were calculated per fold, and folds with fewer than three positives were skipped. Positives were either all test peaks or only the peaks containing at least one pattern_0 instance. Values were then averaged over the five folds. The agreement between perturbed and intact predictions was measured as the Pearson correlation across the test peaks of each fold, averaged over folds. The fraction of the above-chance performance of a model that was retained after a perturbation was defined as

$$\frac{\mathrm{AUROC_{perturbed}} - 0.5}{\mathrm{AUROC_{control}} - 0.5},$$

where $\mathrm{AUROC_{control}}$ is the value for the unperturbed sequences with the observed control track.

Across all 2,282 models, TFs were grouped by the subfamily of the HOCOMOCO v14 motif cluster that contains them, with five additional name-based groups: KLF/SP/MAZ (KLF1–17, SP1–9 and MAZ), FOX, SOX, HOX and CTCF. Archived models were excluded from the family panels and the summary tables. The TFs of non-archived models for which the AUROC on all test peaks decreased to ≤0.75 when pattern_0 instances were replaced by random bases (p0rand) are shown as a circle-packing plot, with circle size proportional to the number of models (184 models, 50 TFs).

#### Binding similarity between experiments across cCREs

To compare the genome-wide binding of the 479 experiments in HepG2 cell-line (Fig. S2), we used the ENCODE registry of candidate cis-regulatory elements (cCREs; Moore et al. 2026). For each experiment, the ChIP-seq fold-change bigWig file was downloaded from ENCODE, and the maximum fold change within each cCRE was recorded; cCREs on chromosomes absent from the bigWig file were set to 0. Similarity between two experiments was calculated as the Pearson correlation coefficient of these values over all cCREs.

Peak-set features were calculated from the ENCODE peak file of each experiment. Each peak was represented by a 200-bp window centered on its summit, and only standard chromosomes were retained. For motif enrichment, we used the representative motif of each HOCOMOCO v14 motif-cluster subfamily, trimmed of low-information edge positions (minimum information content 0.3 bits). Motif matches ($P < 5 \times 10^{-4}$) were counted in the windows of the 1,000 top-scoring peaks and in dinucleotide-shuffled copies of these windows. Enrichment was tested using a one-sided Fisher's exact test, and $P$ values were adjusted for multiple testing using the Benjamini–Hochberg procedure. For genomic annotation, peaks were annotated with ChIPseeker (Yu et al. 2015) against UCSC knownGene (hg38), with promoters defined as −2,000 to +200 bp around the transcription start site (TSS). Promoters overlapping a UCSC CpG island were labeled as CGI promoters, and first introns were reported separately.

TFs were grouped by the HOCOMOCO v14 subfamily of their representative motif, and only subfamilies with more than two within-subfamily pairs were retained. For each TF, the mean and the 10th–90th percentiles of its correlations with TFs of the same subfamily and with TFs of other subfamilies are shown, all within HepG2. TFs and subfamilies are ordered by the mean within-subfamily correlation. The remaining panels show the logo of the most informative representative motif of each subfamily, its enrichment in the peaks of each TF, and the genomic-annotation fractions.

#### Epistasis on a linear scale

The analyses in Figures 2 and 3 measure departures from additivity in log counts, i.e., relative to a multiplicative expectation. To test whether the conclusions depend on this choice (Fig. S3), the same knockout predictions were converted to counts, $C = \exp(P) - 1$, and the departure from additivity was expressed as a fraction of the predicted counts of the intact peak. For direct and other motifs, this was calculated as

$$\frac{(C_{\mathrm{wt}} - C_{\mathrm{none}}) - ((C_{\mathrm{direct}} - C_{\mathrm{none}}) + (C_{\mathrm{other}} - C_{\mathrm{none}}))}{C_{\mathrm{wt}}}$$

over peaks containing both direct and other motifs, and for direct sites as

$$\frac{C_{\mathrm{KO,all}} + \sum_i \left(C_{\mathrm{ref}} - C_{\mathrm{KO},i}\right) - C_{\mathrm{ref}}}{C_{\mathrm{ref}}}$$

in the native peak, over peaks containing at least two (merged) direct sites. For each experiment, we calculated the fraction of peaks with values $\geq t$ (synergy) or $\leq -t$ (redundancy), for $t$ from 0 to 2 in steps of 0.02, and then averaged per TF, as for Figures 2 and 3.
